# Impacts of dietary exposure to pesticides on faecal microbiome metabolism in adult twins

**DOI:** 10.1101/2021.06.16.448511

**Authors:** Robin Mesnage, Ruth C E Bowyer, Souleiman El Balkhi, Franck Saint-Marcoux, Arnaud Gardere, Quinten Raymond Ducarmon, Anoecim Robecca Geelen, Romy Daniëlle Zwittink, Dimitris Tsoukalas, Evangelia Sarandi, Efstathia I. Paramera, Timothy Spector, Claire J Steves, Michael N Antoniou

## Abstract

Concerns have been raised as to whether the consumption of foodstuffs contaminated with pesticides can contribute to the development of chronic human diseases by affecting microbial community function in the gut. We provide the first associations between urinary pesticide excretion and the composition and function of the faecal microbiome in 65 twin pairs in the UK. Biomonitoring of exposure to 186 common insecticide, herbicide, or fungicide residues showed the presence of pyrethroid and/or organophosphorus insecticide residues in all urine samples, while the herbicide glyphosate was found in 45% of individuals. Other pesticides such as DEET, imidacloprid or dithiocarbamate fungicides were less frequently detected. While the geographic location or the rural/urban environment had no influence on pesticide urinary excretion, food frequency questionnaires showed that DMTP levels, a metabolite of organophosphates, was higher with increased consumption of fruit and vegetables. Multivariable association between urinary pesticide excretion and faecal microbial composition and function were determined with shotgun metagenomics and metabolomics. A total of 34 associations between pesticide residues concentrations and faecal metabolite concentrations were detected. Glyphosate excretion was positively associated to an increased bacterial species richness, as well as to fatty acid metabolites and phosphate levels. The insecticide metabolite Br2CA, reflecting deltamethrin exposure, was positively associated with the mammalian phytoestrogens enterodiol and enterolactone, and negatively associated with some N-methyl amino acids. Urine metabolomics performed on a subset of samples did not reveal associations with the excretion of pesticide residues. Our results highlight the need for future interventional studies to understand effects of pesticide exposure on the gut microbiome and possible health consequences.

## Introduction

Human exposure to pesticides have been linked to a variety of diseases triggered by acute intoxication (Boedeker et al. 2020), occupational exposures or residential proximity to pesticide applications (Gonzalez-Alzaga et al. 2015; Gunier et al. 2017; von Ehrenstein et al. 2019). Whether typical low levels of pesticide exposure stemming from dietary and domestic use can contribute to disease development is strongly debated. Nevertheless, adverse effects from chronic exposure during vulnerable periods like pregnancy are well known for some insecticides, such as organophosphates (Bouchard Maryse et al. 2011; Ntantu Nkinsa et al. 2020; Rauh et al. 2011), DDT (Cohn et al. 2015) and pyrethroids (Quirós-Alcalá et al. 2014; Viel et al. 2017).

Controversies around human health effects of pesticides largely originate from the limited ability of current risk assessment procedures employed by government regulatory agencies to predict chronic adverse effects. Animal model systems have been traditionally used to evaluate the toxicity of pesticides. However, toxic effects of pesticides are not always accurately detected in the battery of animal bioassays performed during precommercial stages of assessment. This is the case for neurodevelopmental toxic effects (Fritsche et al. 2018), cancer caused by early life exposures (Manservisi et al. 2017), as well as metabolic disorders and fertility problems caused by endocrine disruptors (La Merrill et al. 2020). This also holds true for the consequences of pesticide exposure on the gut microbiota (Tsiaoussis et al. 2019), which are of interest because of the large enzymatic repertoire harboured by gut microorganisms conferring them the ability to modify the toxicity of chemicals. In some cases, the toxicity of xenobiotics can be enhanced after direct chemical modification by the gut microbiome (Koppel et al. 2017). This has been linked to a variety of health outcomes locally in the gut, such as intestinal damage and severe diarrhea (Wallace et al. 2010), but also at distant organ sites, such as for melamine-induced renal toxicity (Zheng et al. 2013). Xenobiotics can also affect human health indirectly by changing gut microbiome composition (Dethlefsen et al. 2008), decreasing the protective effects of some bacteria or modulating the production of bacterial metabolites (Clayton et al. 2009; Sun et al. 2018).

Dietary habits have a profound influence on the metabolic activity of gut microbial species and their influence on human metabolism. Healthy dietary patterns strongly associate with gut microbiota profiles known to be cardiometabolic markers of health (Asnicar et al. 2021). The gut microbiome responds rapidly to dietary changes (David et al. 2014). Switching to an animal-based diet cause an increase in bile acid secretion to cope with the higher fat intake and this selects for bacteria that are resistant to bile acid. By contrast, switching to a plant-based diet favour bacteria which can use plant polysaccharides. This modulates the capacity of gut bacteria to synthesize vitamins and cofactors (De Angelis et al. 2020). Although the effects of varying levels of macronutrients on the metabolism of gut bacteria are increasingly studied, little is known about the effects of possible contaminants such as pesticides.

Estimating population-level exposure to pesticide residues by direct biomonitoring is becoming one of the most successful strategies to evaluate potential chronic health effects in humans. Analytical chemistry methods sufficiently sensitive to accurately quantify environmental levels of exposure to toxicants in a short turnaround time have been developed in the last decade (Robin et al. 2020). Non-targeted metabolomics technologies are sufficiently sensitive to estimate the levels of food components (e.g. caffeate, curcumin, cinnamate, rosmarinate or daidzein) and a large range of drugs (e.g. metformin, fluoxetine, omeprazole or candesartan) in biospecimen (Visconti et al. 2019). Since the level of pesticides is generally several orders of magnitude lower than endogenous substances, food chemicals, or drugs, detecting these requires more specific assays (Rappaport et al. 2014). Recent initiatives have been launched to harmonise and aggregate pesticide biomonitoring data in the EU with the European Joint Program HBM4EU (Apel et al. 2020), in the US with the CDC’s National Health and Nutrition Examination Survey (Calafat 2012), or the French national programmes Elfe (French Longitudinal Study since Childhood) and Esteban (Environment, Health, Biomonitoring, physical Activity, Nutrition) (Dereumeaux et al. 2017).

No comprehensive biomonitoring of pesticide exposure has been undertaken to date in the UK population. The first aim of our project is to start to fill this crucial gap in our knowledge by studying the exposure to pesticides in 65 twin pairs in the UK. Exposome-wide association studies (ExpWAS) are increasingly performed to link chemical exposures with human disease development (Vineis et al. 2017). This strategy allowed linking exposures between the pesticide-derivative heptachlor epoxide and Type 2 Diabetes (Patel et al. 2010). We provide the first associations between urinary pesticide excretion and the composition and function of the faecal microbiome after deep molecular phenotyping. For this purpose, the faecal microbiome of the 65 twin pairs was studied by shotgun metagenomics and metabolomics to allow associations to be made between dietary factors, pesticide exposure, and faecal microbiome composition and function. Since changes in gut metabolism assessed by analysis of the faecal microbiome can occur without consequences on the relative abundance of the bacterial community, the combination of metagenomics and metabolomics has proven to be the method of choice to study the faecal metabolic environment (Visconti et al. 2019), and to evaluate the disturbance of this ecosystem by pesticides (Mesnage et al. 2021). In addition, we performed a targeted urine metabolomics analysis in order to evaluate whether pesticide urinary excretion also associates with physiological changes (Tsoukalas et al. 2017).

Our study provides the first comprehensive overview of human exposure to pesticides in the UK human population by direct biomonitoring, revealing a widespread exposure to different insecticide residues while the contamination by fungicides and herbicides was less frequent. Analysis of dietary choices further suggested that insecticide exposure was due to the ingestion of contaminated fruit and vegetables. Associations between pesticide excretion and faecal microbiome composition were detected, suggesting that pesticides can be metabolised by gut bacteria although health implications remained unclear. Overall, our study lays the foundation for larger ExpWAS studies linking the exposome to human health outcome by deep molecular phenotyping.

## Results

### Pesticide exposure in the UK population

We first screened urine samples for the presence of 186 pesticide residues in a highly multiplexed detection assay with a low detection limit of 0.1 μg/L per compound (Table S2). This included residues from common insecticides, herbicides and fungicides, which are used in agricultural and domestic settings. When a pesticide was detected, it was included in a follow-up targeted assay to accurately quantify urinary concentrations against a standard curve. Insecticides were the most commonly detected pesticides in all urine samples (Table 1). Pyrethroid and organophosphorus residues were the most abundant, followed by DEET and imidacloprid. The only herbicide detected was glyphosate (Figure 1), which was found in 53% of the urine samples although below the LOQ (< 0.1 μg/L) in 10 cases (8%). Glyphosate was below the limit of detection (< 0.05 μg/L) in the 58 remaining samples analysed (47%). The metabolite of glyphosate AMPA was only quantifiable in three samples (maximum 1.4 μg/L), while it was below the LOQ (< 0.5 μg/L) in seven samples. Exposure to dithiocarbamate fungicides was reflected by the detection of carbon disulphide in 10.8% of the samples.

**Table 1.** Summary statistics. The summary statistics values (μg/L) were estimated using the maximum likelihood inference for left-censored values. <LOD indicates that a reliable value could not be estimated because less than 50% of the samples contained quantifiable amounts of a given compound. In addition to these compounds, fipronil sulfone was detected in 1 sample (LOD of 0.1μg/L) but not quantified. * 6 samples were missing. LOD, limit of detection; DF, detection frequency; IQR, 5 percent and 95 percent quantiles. Pesticide classification is based on information from the European Human Biomonitoring Initiative (HBM4EU).

| pesticide group | active ingredient | metabolite | LOD | DF | median | max | IQR |
| --- | --- | --- | --- | --- | --- | --- | --- |
| dithiocarbamates | dithiocarbamates | carbon disulphide (CS<sub>2</sub>) | 5 | 10.8 | <LOD | 83 | <LOD |
| pyrethroids | cypermethrin, permethrin, cyfluthrin | trans 2,2-dichlorovinyl-2,2-dimethylcyclopropane-1-carboxylic acid (Trans Cl<sub>2</sub>CA) | 0.02 | 96.9 | 0.18 | 23.0 | 0.027 – 1.2 |
|  |  | cis 2,2-dichlorovinyl-2,2-dimethylcyclopropane-1-carboxylic acid (Cis-Cl<sub>2</sub>CA) | 0.01 | 98.4 | 0.07 | 7.1 | 0.014 – 0.38 |
|  | most pyrethroids | 3-phenoxybenzoic acid (3-PBA) | 0.015 | 80.0 | 0.12 | 10.6 | 0.039 – 1.8 |
|  | cyfluthrin | 4-Fluoro-3-phenoxybenzoic acid (4F-3-PBA) | 0.015 | 10.0 | <LOD | 0.10 | <LOD |
|  | deltamethrin | cis-3-(2,2-dibromovinyl)-2,2-dimethylcyclopropane-1-carboxylic acid (Br<sub>2</sub>CA) | 0.015 | 95.4 | 0.077 | 1.64 | 0.014 – 0.42 |
| organophosphorus | Unspecific of methyl-organophosphates, e.g., dimethoate, chlorpyrifos-methyl, azinphos-methyl, malathion, fenthion, phosmet | dimethylphosphate (DMP) | 1 | 14.6 | <LOD | 51.6 | <LOD |
|  |  | dimethylthiophosphate (DMTP) | 1 | 58.5 | 5.2 | 88.6 | 0.83 – 31.9 |
|  |  | dimethyldithiophosphate (DMDTP) | 1 | 2.3 | <LOD | 36.5 | <LOD |
|  | Unspecific metabolite of ethyl-organophosphates e.g., chlorpyrifos, diazinon, ethion, coumaphos, terbufos | diethylphosphate (DEP) | 0.5 | 75.4 | 2.5 | 180 | 0.35 – 17.4 |
|  |  | diethylthiophosphate (DETP) | 1 | 1.5 | <LOD | 5.7 | <LOD |
| glyphosate | glyphosate | glyphosate* | 0.05 | 53 | 0.045 | 2.8 | 0.0025 – 0.84 |
|  |  | aminomethylphosphonic acid (AMPA)* | 0.1 | 5.6 | <LOD | 1.4 | <LOD |
| N,N-dialkylarylamides | DEET | N,N-Diethyl-meta-toluamide (DEET)* | 0.1 | 11.3 | <LOD | 8 | <LOD |
| neonicotinoid | imidacloprid | imidacloprid | 0.05 | 1.6 | <LOD | 1.1 | <LOD |

**Figure 1.**
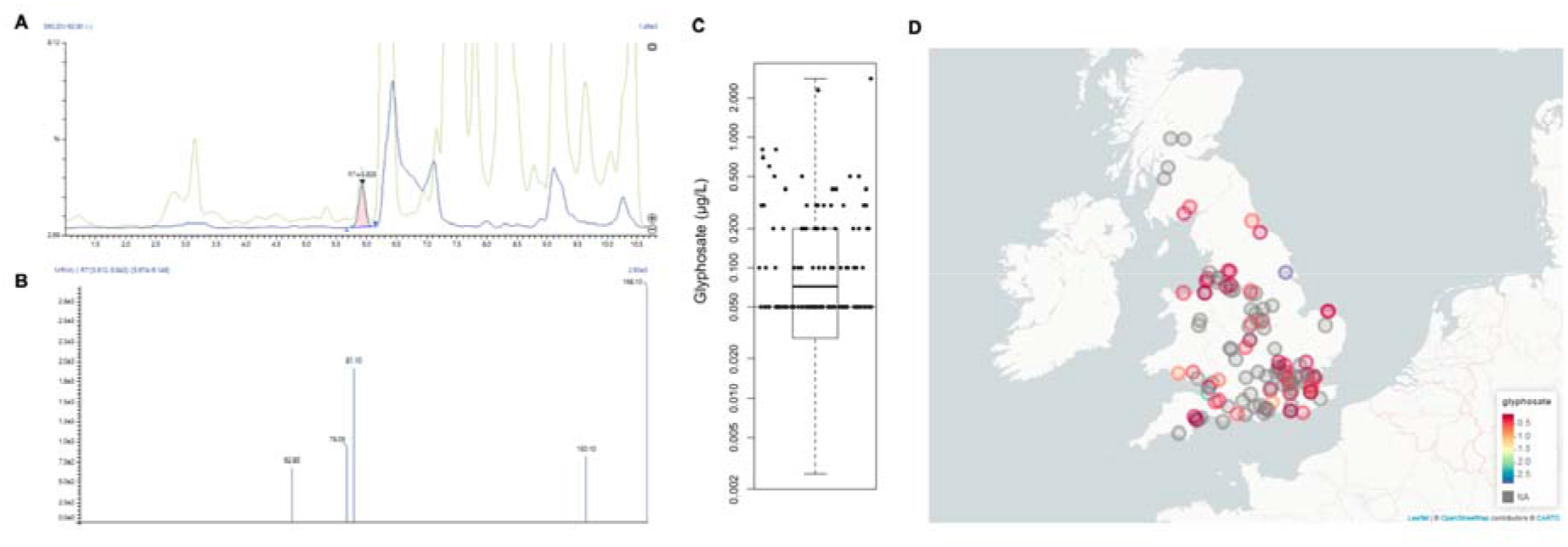
Urinary concentrations of glyphosate in 124 individuals. **A.** Detection of glyphosate in a urine sample spiked with 0.1 μg/L of glyphosate (the limit of quantification in our study). **B.** MRM transition spectrum for the same sample. **C.** Urinary glyphosate levels as a boxplot with the highest censoring threshold (LOQ) shown as a horizontal line. Individual values are added as jittered dots. **D.** Glyphosate levels according to living areas.

### Role of the diet as a determinant of pesticide exposure

We evaluated if dietary habits are associated with the excretion of pesticide residues. This study was originally designed to assess if regular consumption of organic food products results in different urinary pesticide levels. A total of 65 pairs of monozygotic twins discordant for organic food consumption (one twin eats an organic diet whereas the other does not) were selected for this investigation. When a new FFQ was performed at the time of urine collection on these 65 twins, only 15 pairs were discordant for organic food consumption. Although there was no difference in pesticide excretion between twin pairs who reported consuming organic food and those who did not (Figure S1), the inconsistency in the answers provided to the nutrition questionnaire, an issue raised in other studies (Tollosa et al. 2017), convinced us to drop this component of the investigation, as any findings would be deemed as inconclusive.

The role of diet quality in influencing pesticide exposure was then studied using the Healthy Eating Index 2010 (HEI) and a pesticide exposure index created from the consumption of fruits and vegetables (Bowyer et al. 2018; Guenther et al. 2013). Regression between the maximum likelihood estimation of urinary pesticide levels and fruits and vegetable consumption suggested that the exposure to DMTP levels, a metabolite of methyl-organophosphates, is related with the consumption of these types of foods (p = 0.04). The overall evaluation of the model was significant (p = 0.00091). DMTP levels were also associated with the pesticide exposure index created from the consumption of fruits and vegetables (p = 0.01, Figure S2). Although the relationship between pesticide levels and the HEI were not statistically significant, the trends were comparable (Figure S2). This suggested that organophosphate exposure is at least in part related to food contamination.

We also analysed if geographic location could predict urinary pesticide levels (Figure S3). Kruskal-Wallis rank sum test comparing urinary pesticide excretion across 9 UK regions was not statistically significant (see example of glyphosate, Figure 1D). Postcode was available for 123 individuals, which were stratified as 30 rural and 93 urban individuals. Similarly, we did not find a difference in pesticide excretion between rural or urban individuals (Figure S4).

### Deep phenotyping of the faecal microbiome

Faecal metabolite profiles contained xenobiotics, including 84 food components and 46 compounds annotated as pharmaceuticals or pharmaceutical metabolites, and a large number of endogenous compounds such as 197 amino acid derivatives, 30 carbohydrates, and 47 cofactors and vitamins, as well as hundreds of lipids, steroids, corticosteroids, and endocannabinoids (Figure 2A). Faecal metabolomics can also allow the study of the exposome. For instance, our detection of some benzoate metabolites (Figure 2A) could indicate exposure to pyrethroid insecticides such as 3,5-dihydroxybenzoic acid (Hu et al. 2019) and 4-hydroxybenzoate (Pankaj et al. 2016) as these are known to be microbial degradation products of these compounds. However, these are not specific as 3,5-dihydroxybenzoic acid is also considered to be a biomarker for the consumption of whole grain(Wierzbicka et al. 2017).

**Figure 2.**
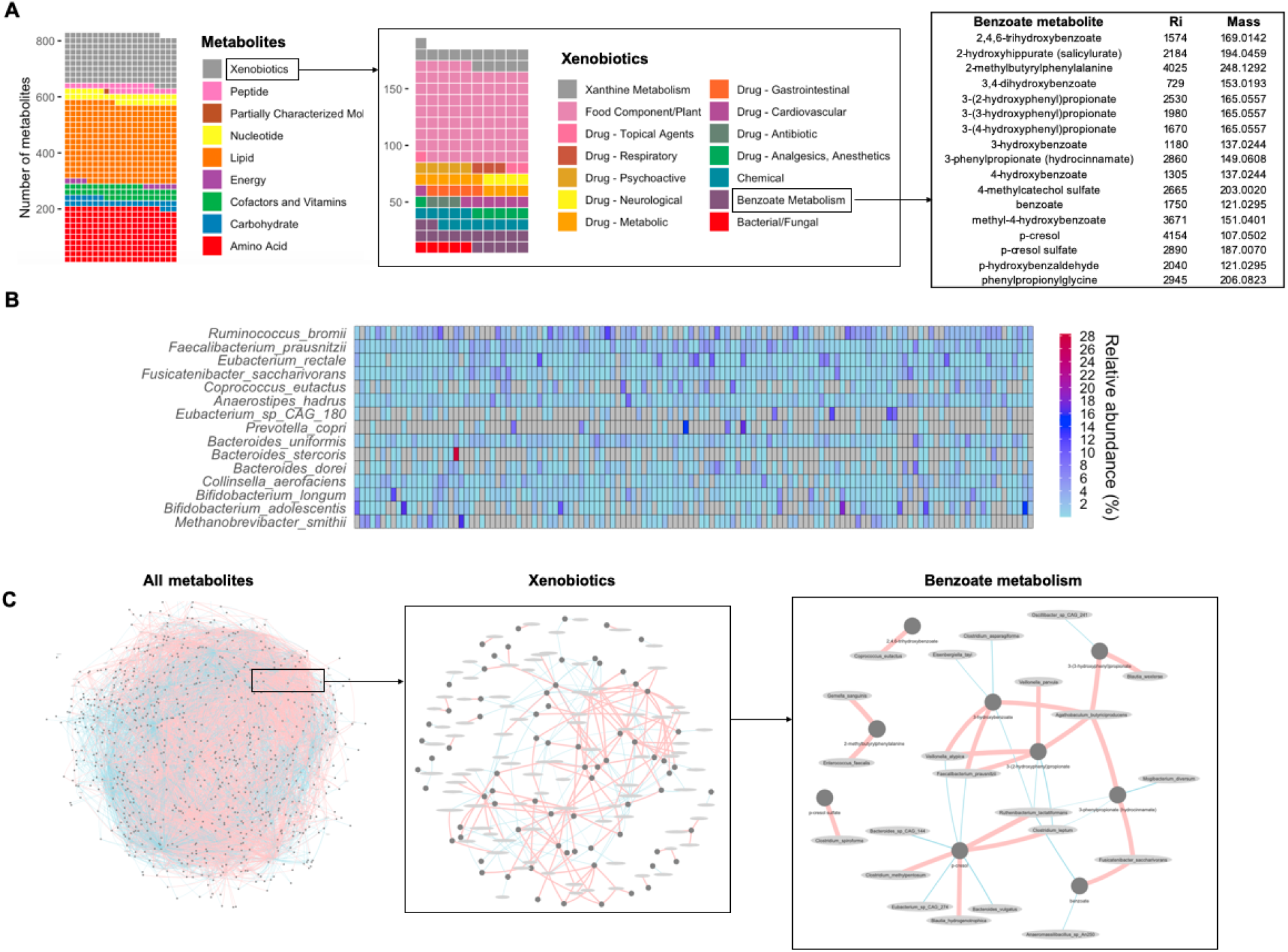
Deep phenotyping of the faecal microbiome in 130 individuals (65 twin pairs). **A**. Faecal metabolomics allow the detection of a large number of metabolites from different classes, including xenobiotics. Taxonomic composition of the gut microbiome was evaluated up to the species level **(B)**. Correlations between the abundance of gut microorganisms and metabolites (positive, red; negative, blue) inform on the interaction between environmental exposures and the metabolism of the gut microbiome.

Taxonomic composition of the faecal microbiome was evaluated by counting clade-specific marker genes. Composition profiles of the faecal microbiomes were typical of faecal microbiome samples from Western developed countries, with the most represented taxa assigned to the phyla Firmicutes (21%) and Bacteroidetes (8%) although the majority of the microorganisms were unknown (64%) and could not be classified by Metaphlan. This large proportion of unclassified microorganisms if not abnormal. It is due to the use of a recent function of Metaphlan allowing the estimation of metagenome composed by unknown microbes which was not included in the calculation of total relative abundance in previous studies using other version of Metaphlan. In total, we identified 16 phyla including 81 families, 205 genera, and 724 species. This were mostly bacteria (637 species), and a few Archaea (5 species), eukaryotes (3 species) and viruses (79 species). Faecal microbiome composition showed high interindividual variation, as only 15 species (out of 21 with an average relative abundance over 1%) were present in 80% of the samples (Figure 2B).

Information on both taxonomic and metabolite composition allows the study of faecal microbiome metabolism (Figure 2C). We detected 8676 correlations between 721 faecal metabolites and 210 bacterial species (FDR < 0.2). Filtering out poorly correlated variables (ρ < 0.3) retained 1949 correlations. There were 148 negative and 144 positive correlations to metabolites classified xenobiotics. These can inform on the interaction between environmental exposures and the metabolism of the gut microbiome. For instance N-(2-furoyl)glycine, which is a furan derivative known to originate from food prepared by strong heating, correlates with multiple species such as *Eggerthella lenta* (ρ = −0.33), *Clostridium bolteae* (ρ = −0.3), *Clostridium citroniae* (ρ = −0.38), *Flavonifractor plautii* (ρ = −0.32), *Methanobrevibacter smithi* (ρ = 0.39). Our data also reveal information about the fate of dietary bioactive compounds, such as the degradation of xanthine metabolites. For instance, *Firmicutes bacterium CAG:110* was negatively correlated to 7-methylurate (ρ = −0.33), paraxanthine (ρ = −0.33), 3-methylxanthine (ρ = −0.37), 1-methylxanthine (ρ = −0.39) and 1-methylurate (ρ = −0.38). Another example is the abundance of saccharine which was correlated with positively correlated with the abundance of *Veillonella dispar* (ρ = 0.39) and *Veillonella infantium* (ρ = 0.38). However, compounds known to be used as pesticides were not detectable by faecal metabolomics.

### Urinary pesticide excretion associates with faecal microbiome metabolism

We evaluated the association between the excretion of pesticides and the composition of the faecal microbiome using MaAsLin2 in order to take into account clinical covariates. Since adjustment by urinary creatinine concentrations can also introduce variability which is not related to urine dilution (Bradman et al. 2013), we also performed the same models without adjustment for creatinine concentrations. A total of 34 associations between urinary insecticide residue concentrations and faecal metabolite concentrations had a FDR below 0.2 when the model was adjusted for creatinine excretion (Table 2). The insecticide metabolite Br2CA, reflecting deltamethrin exposure, was positively associated with the mammalian phytoestrogens enterodiol and enterolactone, as well as negatively associated with some N-methyl amino acids (N-methylalanine, N-methylglutamate, N-2-methylarginine and N-acetyl-1-methylhistidine) (Table 2). Interestingly, faecal 4-hydroxybenzoate, a microbial degradation product of cypermethrin (Pankaj et al. 2016), was negatively associated with trans.Cl2CA concentrations (Table 2). Only associations between Br2CA and N-succinylisoleucine, trans.Cl2CA and oleoyl-linoleoyl-glycerol, as well as trans. Cl2CA and oleoyl-glycerophosphocholine were significant both in unadjusted and adjusted models using insecticide urinary excretion (Table S4).

**Table 2.** Significant associations between urinary excretion of pesticide residues and the composition of the faecal microbiota evaluated using shotgun metagenomics and metabolomics. Statistical models were established with MaAsLin2, using the pesticide levels as predictors. The model coefficient value (effect size) and the standard error from the model are reported along the p-values and its False Discovery Rate (FDR). Associations with FDR < 0.2 for creatinine adjusted models are reported.

| Response | Predictor | effect size | stderr | pval | FDR |
| --- | --- | --- | --- | --- | --- |
|  | **METABOLOMICS** |  |  |  |  |
| Br2CA | N-methylglutamate | -0.15 | 0.04 | 0.0009 | 0.12 |
|  | enterolactone | 0.17 | 0.05 | 0.0011 | 0.13 |
|  | N-succinyl-isoleucine | -0.12 | 0.04 | 0.0012 | 0.13 |
|  | palmitoyl-ethanolamide | -0.18 | 0.05 | 0.0017 | 0.13 |
|  | N-methylalanine | -0.20 | 0.06 | 0.0019 | 0.13 |
|  | 2-hydroxyglutarate | -0.16 | 0.05 | 0.0026 | 0.14 |
|  | N-acetyl-1-methylhistidine. | -0.18 | 0.06 | 0.0026 | 0.14 |
|  | enterodiol | 0.19 | 0.06 | 0.0030 | 0.14 |
|  | N2-methylarginine | -0.07 | 0.02 | 0.0038 | 0.16 |
|  | piperidine | -0.15 | 0.05 | 0.0041 | 0.16 |
|  | stearoyl-ethanolamide | -0.21 | 0.08 | 0.0060 | 0.17 |
|  | 2-oxo-1-pyrrolidinepropionate | -0.08 | 0.03 | 0.0061 | 0.17 |
|  | dihydroferulate | -0.12 | 0.04 | 0.0073 | 0.19 |
|  | 1-palmitoyl-2-arachidonoyl-GPC | 0.11 | 0.04 | 0.0076 | 0.19 |
| Trans.Cl2CA | oleoyl.linoleoyl.glycerol | 0.23 | 0.07 | 0.0013 | 0.15 |
|  | linoleoyl-linoleoyl-glycerol | 0.23 | 0.07 | 0.0014 | 0.15 |
|  | 4-hydroxybenzoate | -0.14 | 0.05 | 0.0020 | 0.15 |
|  | glycerol | 0.25 | 0.08 | 0.0023 | 0.15 |
|  | 1-oleoyl-GPC18 | 0.25 | 0.08 | 0.0031 | 0.19 |
|  | anacardic.acid | 0.30 | 0.10 | 0.0034 | 0.19 |
| 3.PBA | arachidoylecarnitine | -0.25 | 0.07 | 0.0005 | 0.12 |
|  | carotenediol | -0.13 | 0.04 | 0.0006 | 0.12 |
|  | 3-methylurate | 0.18 | 0.05 | 0.0007 | 0.12 |
|  | allantoin | -0.27 | 0.08 | 0.0007 | 0.12 |
|  | erucoylecarnitine | -0.23 | 0.07 | 0.0008 | 0.12 |
|  | lysine | -0.11 | 0.04 | 0.0027 | 0.18 |
| glyphosate | 3.hydroxystearate | 0.51 | 0.14 | 0.0003 | 0.19 |
|  | lactobacillic acid | 0.25 | 0.07 | 0.0010 | 0.19 |
|  | stearate | 0.25 | 0.07 | 0.0010 | 0.19 |
|  | phosphate | 0.41 | 0.12 | 0.0011 | 0.19 |
|  | 2-hydroxylignocerate. | 0.33 | 0.10 | 0.0011 | 0.19 |
|  | pentadecanoate | 0.24 | 0.07 | 0.0013 | 0.19 |
|  | 3-carboxyadipate | 0.27 | 0.08 | 0.0019 | 0.19 |
|  | saccharopine | 0.34 | 0.11 | 0.0019 | 0.19 |
|  | **METAGENOMICS** |  |  |  |  |
| glyphosate | *Fusicatenibacter saccharivorans* | -0.28 | 0.09 | 0.002 | 0.19 |
| Cis.Cl2CA | *Parabacteroides johnsonii* | 0.08 | 0.0004 | 3.74E-39 | 1.2E-36 |
|  | *Olsenella scatoligenes* | 0.08 | 0.01 | 1.05E-09 | 2.2E-07 |
| DEP | *Parabacteroides johnsonii* | 0.05 | 0.0002 | 2.52E-37 | 7.9E-35 |
| Trans.Cl2CA | *Olsenella scatoligenes* | 0.08 | 0.01 | 1.23E-06 | 0.0008 |
| 3.PBA | *Olsenella scatoligenes* | 0.09 | 0.02 | 0.0001 | 0.03 |
|  | *Streptococcus cristatus* | -0.01 | 0.003 | 0.0002 | 0.03 |
|  | *Clostridium symbiosum* | 0.09 | 0.02 | 7.28E-05 | 0.03 |

Limited associations were found between urinary pesticide concentrations and taxonomic composition when the model was adjusted or unadjusted for creatinine excretion, respectively. Five associations were significant in both, adjusted and unadjusted, models (Table 2 and S4). *Olsenella scatoligenes and Clostridium symbiosum* were positively associated to the metabolism of pyrethroids while a negative association with the urinary levels of these insecticides was found for *Streptococcus cristatus*. Only *Collinsella stercoris* was positively associated with organophosphates levels. Insecticide levels did not significantly influence microbiome diversity in most cases (Table S5), and only the levels of DEP, a metabolite of ethylorganophosphates, was positively associated to the number of observed species.

We recently published the description of a metabolomic signature for glyphosate effects in the gut microbiome of Sprague-Dawley rats (Mesnage et al. 2021). In order to understand if our results in laboratory animals can be translated to human populations, we evaluated if there was a correlation between glyphosate urinary levels and faecal microbiome composition and function in the TwinsUK cohort. Faecal metabolomes from this twin study contained 10 metabolites that were previously found to have their levels altered in the caecum microbiome of rats exposed to glyphosate, namely hydroxy-N6,N6,N6-trimethyllysine, N-acetylputrescine, valylglycine, glutarate, pimelate, linolenoylcarnitine, carotene diol, 3-dehydroshikimate, 2-isopropylmalate and solanidine (Mesnage et al., 2021). Since glyphosate was undetected in approximately half of the samples (47%), we evaluated the relationship between the presence of glyphosate and the composition of the faecal microbiome using a random forest method. The 10 metabolites known to discriminate glyphosate-exposed rats from unexposed rats did not significantly predict the detection of glyphosate in the 124 individuals with a classification accuracy of 65% (95% CI [0.49, 0.79]) (Figure S5). Classification of the samples using the full set of 736 metabolites (accuracy = 0.56, 95% CI [0.40, 0.71]), or the taxonomic composition profiles (accuracy = 0.53, 95% CI [0.38, 0.69]), was also not accurate. Results of linear-mixed models considering age and creatinine excretion as a covariate, as well as the twin relatedness as a random effect, also provided limited insights. The FDR for the most significant associations between individuals who excreted glyphosate and those for whom glyphosate was undetected was relatively high (Figure 3A). Nonetheless, *Fusicatenibacter saccharivorans* was more abundant in individuals excreting glyphosate (p = 0.002, FDR = 0.19) (Figure 3E).

**Figure 3.**
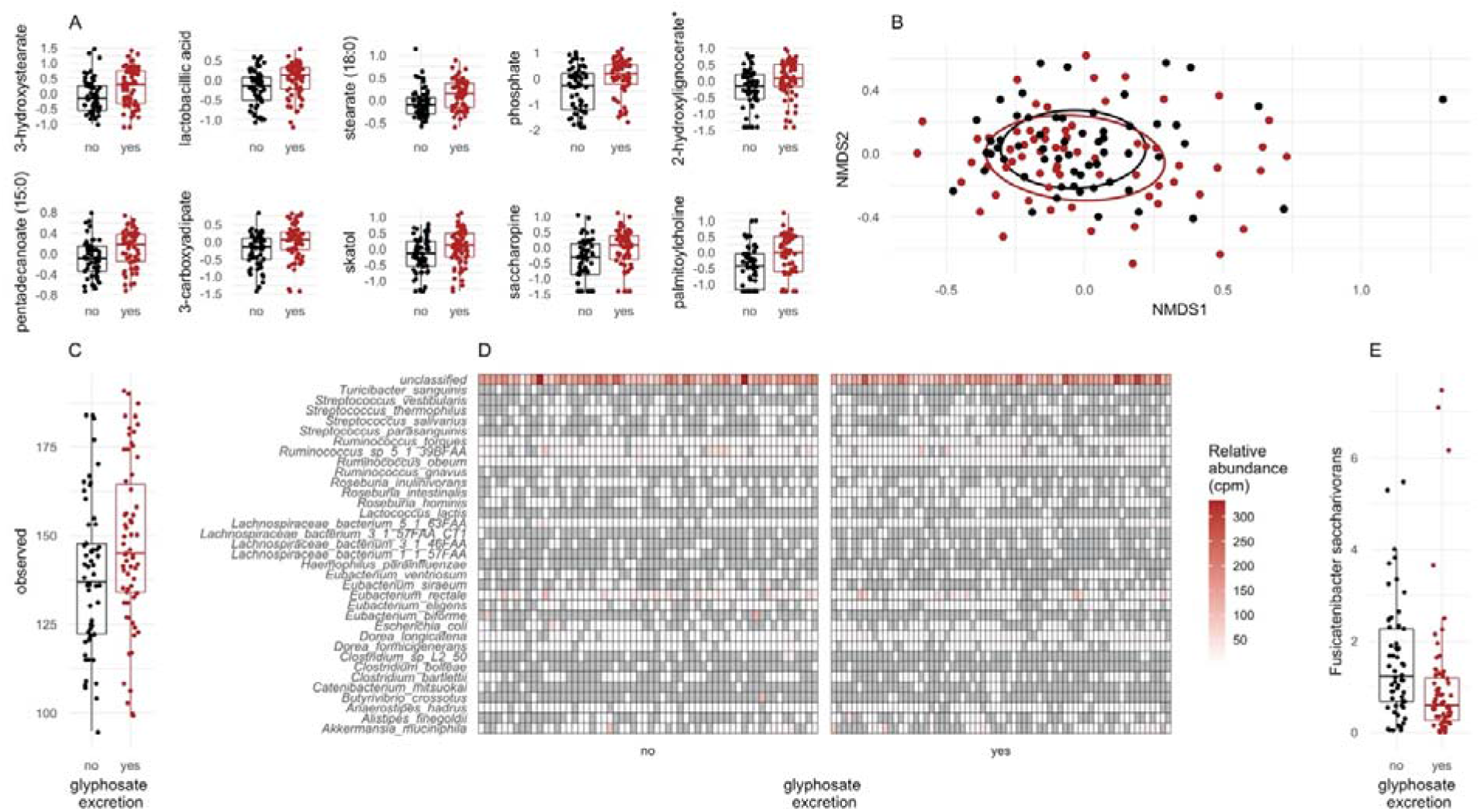
Difference in gut microbiota profiles between individuals who excreted glyphosate and those for whom glyphosate was undetected. **A.** Faecal metabolites, which have the largest difference in abundance. Log-transformed abundance values are shown as box plots. **B**. Beta diversity using Bray-Curtis dissimilarity. **C.** Alpha diversity as the number of observed species. **D.** Relative abundance (as copies per million, cpm) for the bacteria contributing to the abundance of the shikimate pathway (as MetaCyc Pathway: chorismate biosynthesis I). **E.** Difference in relative abundance for *Fusicatenibacter saccharivorans*.

We also estimated the relative contribution of the different bacterial species to core functions of the gut microbiome using Humann2. Although a large number of pathways were identified, the relative contribution of the different bacterial species remained partially characterised, such as in the case of the taxonomy of the shikimate pathway (as MetaCyc Pathway: chorismate biosynthesis I) (Figure 3D). The relative abundance of genes from the shikimate pathway among bacteria, which are identified as major contributors to the total abundance of this pathway, is not different between the individuals who excreted glyphosate and those for whom glyphosate was undetected (Figure S6).

Although glyphosate was not found to inhibit the shikimate pathway using the metabolic signature from our previous laboratory animal experiment (Mesnage et al., 2021), its presence was positively associated with 8 metabolites, some of which could provide insights into its metabolic roles. Positive associations to glyphosate levels mostly included fatty acid metabolites (Table 2). The positive association between phosphate and glyphosate points to an influence on phosphate metabolism, which could be due to the metabolism of glyphosate by the gut microbiome as we have previously hypothesised (Mesnage and Antoniou 2020).

Microbial composition measured by Bray-Curtis dissimilarity was not different between individuals who excreted glyphosate and those for whom glyphosate was undetected (PERMANOVA p= 0.15). A linear mixed model was created to test if alpha diversity could be predicted by the exposure to glyphosate, also adjusting for creatinine, age and total number of DNA sequence reads. The species richness was higher in individuals who excreted glyphosate (p = 0.009) although other indicators of alpha diversity (Shannon and Simpson diversity indices) were not significant.

### Urine metabolome

Urine metabolomics analysis consisted of a targeted analysis of 36 organic acids in a subgroup of 61 subjects. No differences were detected in the concentrations in organic acids between a group of 28 individuals who did not present detectable glyphosate levels in their urine compared to a group of 33 individuals with detectable glyphosate levels (Table S6). No associations between pesticide residues and the composition of the urine metabolome were found using linear-mixed models considering age and creatinine excretion as a covariate, as well as the twin relatedness as a random effect. However, the relatively low number of samples limited the statistical power and probably prevented detection of associations.

## Discussion

The consequences of pesticide exposure on human gut microbial community composition, functions and metabolic health is currently unknown, despite the increasing number of studies in laboratory animals showing perturbations of the gut microbiome by pesticides (Tsiaoussis et al. 2019). We performed the first pesticide biomonitoring survey of the British population, and subsequently used the results of this study to perform the first pesticide association study on gut microbiome composition and function from the TwinsUK registry. The excretion of insecticides was associated with the consumption of fruits and vegetables, as well as to the composition of the faecal microbiome. In addition, although glyphosate was not found to influence the shikimate pathway, its excretion was positively associated to an increased bacterial species richness.

Pesticide exposure at environmental levels have been suggested to disturb gut microbial metabolism. Our results support recent studies suggesting the faecal metabolome is a robust readout of gut microbiome metabolism (Zierer et al. 2018), and that it provides more information than the extrapolation of pathway alterations from microbial gene abundances using metagenomics as we demonstrated recently in laboratory animals exposed to glyphosate (Mesnage et al. 2021). The insecticide metabolite Br2CA, reflecting deltamethrin exposure, was negatively associated with amino acid metabolites (Table 2). Deltamethrin transformation by *Bacillus thuringiensis* has been found to cause a downregulation of energy metabolism (Guo et al. 2020), although a direct comparison of these findings to the gut microenvironment is speculative. Microbial abundance and diversity was not found to be decreased by glyphosate as theorised by some authors (Samsel and Seneff 2013) but increased, which is coherent with the findings of our recent study in rats where we found glyphosate interference with gut microbial metabolism (Mesnage et al. 2021). The positive association between phosphate and glyphosate points to an influence on phosphate metabolism, which could be due to the metabolism of glyphosate by the gut microbiome as hypothesised previously (Mesnage and Antoniou 2020).

To the best of our knowledge, no other studies corroborate the role of *Olsenella scatoligenes, Clostridium symbiosum* or *Streptococcus cristatus* in the metabolism of pyrethroids, or the role of *Collinsella stercoris* in the metabolism of organophosphates. Bacterial species found to have their levels altered in the gut microbiome of laboratory animals exposed to organophosphate insecticides (Roman et al. 2019) were not seen to be altered in this study. This is probably because population exposure levels are substantially lower in comparison to those used in toxicology bioassays. Higher levels of exposure could be necessary to cause changes in microbiome composition, such as in the case of insects becoming resistant to insecticides due to the growth of insecticide-degrading bacteria being selected for in their gut microbiome (Daisley et al. 2018; Kikuchi et al. 2012). Some *Pseudomonas* species even subsist on caffeine as a sole source of energy allowing caffeine detoxification in the primary insect pest of coffee (Ceja-Navarro et al. 2015). Only one other study in a human population has made a direct link between the degradation of organophosphate insecticides in the human gut microbiome and its consequences on glucose metabolism (Velmurugan et al. 2017). However, this study was undertaken among pesticide users who can be exposed to significantly higher levels of pesticides even if they are living in agricultural areas without directly handling pesticides (Deziel et al. 2015). In the case of occupational exposures, individuals are also exposed to the complete commercial formulated products, which could include other ingredients more toxic than the stated active pesticidal compound (Mesnage and Antoniou 2018).

Urinary pesticide levels were comparable to those of previous studies performed on European populations in other countries. Median pyrethroid exposures based on the metabolite 3-PBA have been found to range from 0.018 to 0.43 μg/L in France (Dereumeaux et al. 2018; Viel et al. 2015), Germany (Schulz et al. 2009), Poland (Wielgomas and Piskunowicz 2013), Denmark (Dalsager et al. 2018), and are thus comparable to concentrations from our study (0.12 μg/L). For organophosphates, the median of the molar sum of organophosphates metabolites in our study (34.7 nM, Table S3) was comparable to values reported in Denmark (56.5 nM, n = 564; (Dalsager et al. 2018) and France (38.8 nM, n = 254; (Debost-Legrand et al. 2016). Concerning glyphosate, our results show a relatively high impregnation of the population by glyphosate. In another study on 50 Irish adults, 20% were found to excrete glyphosate (0.80-1.35 μg/L) (Connolly et al. 2018). A German study of 399 urine samples found that 31.8% of the samples contained glyphosate concentrations exceeding a LOQ of 0.1 μg/L (same as our study) (Conrad et al. 2017). Although other glyphosate biomonitoring studies were performed in European populations (Gillezeau et al. 2019), these used an enzyme-linked immunosorbent assay, which has not been validated for urine biomonitoring. Further epidemiological studies will be necessary to ascertain if glyphosate has an impact on the gut microbiome of populations exposed to higher environmental concentrations, such as in the US (Mills et al. 2017) or in the case of occupational exposures (Mesnage et al. 2012).

The consumption of fruit and vegetables is the major source of pesticide exposure in the UK (EFSA 2019). A large range of pesticides were applied in the UK in 2016, including insecticides (316 tonnes), fungicides (5,902 tonnes), herbicides (7,806 tonnes), and molluscicides (161 tonnes)(FERA 2016). The consumption of agricultural products sprayed with pesticides is not always the most important source of exposure as pesticides are frequently found in dust (Colt et al. 2004) and ambient air (Socorro et al. 2016). Pesticides are also used by the amenity sector (e.g., golf courses, local authorities, lawn care operators, sports stadiums), with 80 tonnes of pesticides applied in 2016 (77% by glyphosate, 61 tonnes) (FERA 2016). Domestic use is also an important source of exposure with the use of herbicides to clear weeds in private gardens, or the use of insecticides indoors (e.g. anti-mosquito sprays or impregnated animal pet collars for flea control) (Dyk et al. 2011).

Our results suggest that the exposure to some organophosphate and pyrethroid insecticide residues is related to the diet. This corroborates the results of previous studies, which showed that individuals eating an organic diet had lower levels of urinary dialkylphosphate (DAP) (Curl et al. 2003) and 3-PBA (Curl et al. 2019) than individuals who were eating conventional products. In another investigation, ingestion of 100[grams of fruit increased urinary DAP excretion by 7% (van den Dries et al. 2018). Organophosphorus compounds are widely found in the environment, and human exposure to these chemicals can also come from other sources such as house dust (Colt et al. 2004). The exposure to organophosphate during sensitive periods of life has been linked to a variety of diseases such as neurobehavioral problems after prenatal exposures (Bouchard Maryse et al. 2011; Sagiv et al. 2019) or neurodegenerative diseases like Parkinson’s disease (Wang et al. 2014). Although we could not test whether pesticide excretion is different between individuals eating organic food and others who do not, the observed inconsistency between the results of the FFQ before recruitment and during the study is also an important result, which supports the need for intervention studies (Bradman et al. 2015; Curl et al. 2019). An organic diet is multifactorial and difficult to clearly define, which makes self-assessment prone to subjective biases (Mesnage et al. 2020). New studies would need to focus on vulnerable groups, such as pregnant farmworkers who do not necessarily have access to proper safety equipment in rural environments and are thus exposed to higher pesticide levels (Curl et al. 2020).

The major limitation of this study is its sample size. The number of individuals we investigated is sufficient to provide reliable information on environmental levels of exposure, since reference values in biomonitoring studies require a sample size ranging from 73 to 120 individuals (Vogel et al. 2019). However, our sample size is low to find associations between pesticide excretion and gut microbiome composition. Small differences in alpha diversity (an effect size of 0.55) between two groups of 55 individuals can be detected with 80% statistical power (Casals-Pascual et al. 2020), which provides sufficient power to suggest that the increased microbial diversity observed in individuals excreting detectable levels of glyphosate (Figure 3C) is reliable. However, gut microbiome taxonomic data is typically over-dispersed and zero-inflated (Knight et al. 2012). There is no gold standard for statistical analysis of EWAS data, and it is not clear how a list of statistically significant associations can be translated to information usable for public health policies (Cheung et al. 2020). In our study, more than 50% of the datapoints were equal to 0 for 612 species out of the 724 detected. In this case, when a value is 0, it is not clear whether the species is absent or undetected. In addition, a large number of unidentified factors influence the results of gut microbiome studies (McLaren et al. 2019). These factors can be technical covariates such as the DNA extraction procedure (Costea et al. 2017; Ducarmon et al. 2020), or the sequencing approach (Singer et al. 2019), demographic differences (Rampelli et al. 2015), lifestyle changes such as the intake of prescription medications (Jackson et al. 2018), alcohol consumption frequency and bowel movement quality (Vujkovic-Cvijin et al. 2020), or even socioeconomic factors (Bowyer et al. 2019). In addition, there is no gold standard for hardware and software for taxonomic assignment of shotgun metagenomics datasets (Ye et al. 2019). Our study is thus a first step towards the understanding of pesticide-induced gut microbial changes in human populations, but larger studies will be needed.

In conclusion, this study provides the first evidence of an association between pesticide excretion and changes in gut microbiome metabolism at environmental levels of exposure in the UK population. This was partly attributed to the consumption of fruit and vegetables, which can be sprayed with insecticides for pest control. This highlights the need for future interventional studies to understand the role of the gut microbiome as a mediator of health and pesticide exposure. We demonstrate that molecular signatures characterised using high-throughput ‘omics’ technologies in animal studies can be found in the human population to evaluate perturbation of toxicity pathways. Although more studies are needed to understand health consequences, our study provides a foundation for the development of new environmental epidemiology studies linking pesticide exposure to metabolic perturbations and their health consequences.

## Material and methods

### Participants and pesticide exposure estimation

Subjects were monozygotic twins enrolled in the TwinsUK cohort (Verdi et al. 2019). The St. Thomas’ Hospital Research Ethics Committee approved the study. All individuals provided informed written consent. Twins were selected based on their answers to a food frequency questionnaire (FFQ) modified to include a question on organic food consumption (Bowyer et al. 2018). Our aim was to define two groups of individuals, one less likely to be exposed to pesticides than the other because of organic food consumption. Consumption of legumes, fresh fruits and vegetables were estimated using existing FFQ data following the EPIC-Norfolk guideline (EPIC 2017). Relevant FFQ items were converted to grams consumed per week (for conversion table, please see Bowyer et al 2018.) Responses to the additional question “*Please indicate to what extent you consume, when available, organic fruits and vegetables*?” were used as modifiers to estimate potential for pesticide exposure, with the per weekly gram consumption being multiplied by the relevant weight (Table S1). Individuals who responded that they did no eat fruits and vegetables were removed from analysis. This resulted in a proxy estimate for potential of pesticide exposure from the diet (hereafter referred to as ‘pesticide exposure’). Differential pesticide exposure between twin pairs was assumed for pairs with a >1 standard deviation difference of estimated exposure and who fell within different categories. A total of 977, mostly female, twin pairs answered questions on organic food dietary intake from the TwinsUK questionnaire. Among these 977 individuals, 65 twin monozygotic twin pairs were found to be discordant for organic food consumption.

Geographic location of the individuals from this study was based on their postcode centroid. The 111 individuals with geographic location were from different UK regions, namely East Midlands (9), East of England (16), London (10), North East (2), North West (15), South East (30), South West (19), West Midlands (6), and Yorkshire and The Humber (4). The discrimination between rural and urban environments was established with the Land Cover Map 2015 (LCM, version 1.2), which was downloaded from the Centre for Ecology and Hydrology via the ‘Digimap’ portal. Individuals were considered as rural or urban based on their surrounding environment in a 1 km^2^ area.

### General pesticide screening in urine samples

Urine was extracted with acetonitrile and QuEChERS salt and then injected into the LC/MS-MS system. For this, Albendazol, Carbendazim ^3^D, Phosalone ^10^D were used as internal standards (IS). Albendazol (LGC, UK) at a purity of 98.1 % was dissolved in acetonitrile and 10 % formic acid to obtain a working solution of 2 mg/L. Carbendazim ^3^D (LGC, UK) at a purity of 96.6 %, was dissolved in dimethylformamide to obtain a working solution of 2 mg/L. Phosalone ^10^D purchased as a solution at 100 mg/L (LGC, UK), was diluted in acetonitrile to obtain a working solution of 1 mg/L.

For sample preparation and extraction, 10 μL of IS and 5 mL of acetonitrile was added to 1 mL of urine. The sample was agitated before centrifugation at 3000 g for 5 min and the supernatant was transferred to a 15 mL glass tube. One gram of QuEChERS Mix 1 (MgSO_4_/NaCl/C_6_H_5_Na_3_O_7_, 2H_2_O/C_6_H_6_Na_2_O_7_, 1.5H_2_O) (4/1/1/0.5) (w/w/w/w) was added to the sample and agitated before being centrifuged again at 3000 g for 5 min. The organic phase was then transferred to a 15 mL glass tube and evaporated to dryness under nitrogen flow. The dried sample was taken in 200 μL of (50/50) mobile phase solutions. Only 2 μL of the re-dissolved sample solution was injected into the LC-MS/MS system.

The LC-MS/MS system included a Shimadzu NEXERA X2 series and a 8060 triple quadrupole mass spectrometer. Chromatographic separations were performed on a Raptor Biphenyl column (100 × 2.10 mm, 2.7 μm particles) (Restek, France). Mobile phase A contained 0.002 % formic acid in 2 mM ammonium formiate and phase B consisted of methanol and 0.002 % formic acid in 2 mM ammonium formiate. Identification and quantification of pesticides was performed in positive and negative mode using multiple reaction monitoring (MRM) of a quantification and additional qualifier ion. To meet the criteria for a positive identification, the ratio between the quantitative and the qualifying transition ions had to fall within ±20% of that established by the calibration standards.

### Glyphosate biomonitoring in urine samples

Glyphosate and its major metabolite, aminomethylphosphonic acid (AMPA), were successfully measured in 124 urine samples (note: an insufficient volume of urine was available for the remaining 6 samples). Glyphosate and AMPA were measured after a derivatization reaction using FMOC-Cl (9-fluorenylmethyl chloroformate). Samples were then extracted with diethyl ether and injected into the LC/MS-MS system. Glyphosate ^13^C2^15^N was used as an IS, and was purchased as a solution at 100 mg/L (LGC, UK) and diluted in deionized water to obtain a working solution of 0.5 mg/L. Glyphosate and AMPA (LGC, UK) were of 98.69% and 99% purity respectively, and were dissolved in deionized water to obtain working solutions at increasing concentrations ranging from 0.01 to 50 mg/L. These standard solutions were used to spike glyphosate-free urine for the preparation of the calibration curves for standards. Six calibration standards between the higher limit of quantification (LOQ) and the lower LOQ (namely, between 0.1 to 10 μg/L) were necessary for the calibration.

FMOC (Acros Organics, Belgium) was prepared at 50 g/L and was used for the derivatization reaction. Glyphosate and AMAP already derivatized with FMOC were purchased from LGC (98 %, 99.6 % purity respectively). Working solutions of Glyphosate-FMOC and AMAP-FOMC at 0.1 and 1 mg/L were used to spike glyphosate-free urine samples to prepare internal quality controls at 0.5 and 5 μg/L.

A 50 μL volume of IS and 1 mL of 0.5 M tetraborate buffer (pH 9) were added to 1 mL of urine. Then, 3 mL of the FMOC solution was added and the sample allowed to stand for 30 min in the dark. For the extraction of the formed derivatives, 1 mL of 6M HCl and 6 mL of diethyl ether were added to each sample and agitated for 15 min before centrifugation at 3000 g for 5 min. The organic phase was then transferred to a 15 mL glass tube and evaporated to dryness under nitrogen flow. The dried sample was taken up in 200 μL of (50/50) mobile phase solutions and a 10 μL aliquot injected into the LC-MS/MS system. The calibration standards were treated in the same way after spiking of the appropriate volume of the working solutions.

The LC-MS/MS system included a Shimadzu NEXERA X2 series and a 8060 triple quadrupole mass spectrometer. Chromatographic separations were performed at 40°C on a Kinetex C18 100A column (100 × 2.10 mm, 2.6 μm particles) (Phenomenex, France). Mobile phase A contained 0.05% formic acid and phase B included acetonitrile and 0.05% formic acid. Identification and quantification of glyphosate-FMOC and AMPA-FMOC were performed in negative mode using MRM of a quantifier ion (390.2/62.9 and 331.9/110.1, respectively) and an additional qualifier ion (389.9/168.1 and 331.9/62.9, respectively). To meet the criteria for a positive identification, the ratio between the quantitative and the qualifying transition ions (derived from the precursor ion) had to fall within ±20% of that established by the calibration standards.

### Biomonitoring of pyrethroid metabolites in urine samples

Pyrethroid metabolites are measured in urine after hydrolysis with β-glucuronidase (Helix Pomatia). They are extracted with hexane and injected into the LC/MS-MS system.

For this, 3-PBA ^13^C and trans-Cl_2_CA ^6^D were used as IS. Both IS, purchased as solution at 100 mg/L (LGC, UK and CLUZEAU INFO LABO, France), were diluted in (50/50) acetonitrile and 2mM ammonium formiate buffer (pH 3) at 0.05 mg/L to obtain IS working solutions. For calibration curves, working solutions containing 3-PBA in methanol at 1 g/L, 4-FPBA at 100 mg/L in acetonitrile, 2,2-dichlorovinyl-2,2-dimethylcyclopropane-1-carboxylic acid (Cl_2_CA) (cis and trans) at 10 mg/L in methanol (CLUZEAU INFO LABO, France) and cis−3-(2,2-dibromovinyl)-2,2-dimethylcyclopropane-1-carboxylic acid (Br_2_CA) at 10 mg/L in methanol (CLUZEAU INFO LABO, France) were prepared at increasing concentrations ranging from 0.01 to 2 mg/L. These solutions were used to spike pyrethroid-free urine for the preparation of the calibration curve standards. Six calibration standards between the higher LOQ and the lower LOQ (namely, between 0.025 to 10 μg/L) were necessary for the calibration.

To 5 mL of urine, 25 μL of IS and 1.25 mL of 1 M sodium acetate buffer (pH 4.8) were added. Then, 20 μL of β-glucuronidase (Helix Pomatia) was added and samples incubated overnight for 16 h at 37 °C. For the extraction of the hydrolyzed molecules, 1 mL of 37 % HCl and 6 mL of hexane were added to the samples and agitated for 10 min before centrifugation at 2000 g for 5 min. The organic phase was then transferred to a 15 mL glass tube. Hexane (6 mL) was added to the remaining aqueous phase and agitated for 10 min before centrifugation at 2000 g for 5 min. The two organic phases were then combined into one tube. Then, 3 mL of 0.1 M NaOH was added to this final organic phase and agitated for 10 min before centrifugation at 2000 g for 5 min. After removal of the upper phase, 200 μL of 37 % HCl and 6 mL of hexane were added to the samples and agitated for 10 min before centrifugation at 2000 g for 5 min. The upper phase was then transferred to a 10 mL glass tube and evaporated to dryness under nitrogen flow. The dried sample was taken up in 80 μL of water and 0.1 % formic acid (70/30) and 10 μL injected into the LC-MS/MS system. The calibration standards were treated in the same way after spiking of the appropriate volume of the working solutions.

The LC-MS/MS system included a Shimadzu LC-20AD and AB SCIEX API 5500 QTrap triple quadrupole mass spectrometer. Chromatographic separations were performed on a Atlantis T3 column (150 × 2.10 mm, 5 μm particles) (Waters, USA). Mobile phase A contained 0.1% formic acid and phase B included (95/5) methanol acidified with 0.1% formic acid and phase A. Identification and quantification of 3-PBA, 4-FPBA, Cl_2_CA (cis and trans) and Br_2_CA were performed in negative mode using MRM of a quantifier ion (213.0/92.9, 231.0/93.1, 208.9/36.9 and 342.9/80.8, respectively) and an additional qualifier ion (213.0/65.1, 231.0/65.1, 207.0/35.0 and 296.8/80.9, respectively). To meet the criteria for a positive identification, the ratio between the quantitative and the qualifying transition ions (derived from the precursor ion) had to fall within ±20% of that established by the calibration standards.

### Organophosphate metabolite biomonitoring

Organophosphate metabolites (dialkyl phosphate, DAP) are measured in urine after an extraction with ethyl acetate and diethyl ether. They are then injected into the LC/MS-MS system. DMP ^6^D, DMTP ^6^D, DMDTP ^6^D, DEP ^10^D, DETP ^10^D and DEDTP ^13^C_4_ were used as IS. DMTP ^6^D, DMDTP ^6^D and DEDTP ^13^C_4_ (LGC, UK) at 97 %, 98 % and 95 % purity respectively, were dissolved in methanol to obtain a working solutions of 1 g/L. DMP ^6^D and DEP ^10^D (CLUZEAU INFO LABO, France) at purities of 95 % and 99 % respectively, were dissolved in methanol to obtain working solutions of 1 g/L. DETP ^10^D was purchased as a solution at 100 mg/L (CLUZEAU INFO LABO, France). IS solutions were mixed to obtain a working solution of 1 mg/L per IS. DMP, DETP and DEDTP (SIGMA-ALDRICH, USA) at purities of 100 %, 98 % and 90 % respectively, were dissolved in methanol. DMTP and DMDTP (LGC, UK) of 96 % and 95 % purity respectively, were dissolved in methanol. DEP (CHEM Service, USA) at a purity of 99.5 % was dissolved in methanol. DMP, DMTP, DMDTP, DEP, DETP and DEDTP were mixed to obtain working solutions at increasing concentration ranging from 0.1 to 10 mg/L. These standards solutions were used to spike DAP-free urine for the preparation of the calibration curve standards. A total of 6 calibration standards between the higher LOQ and the lower LOQ (namely, between 2 to 100 μg/L) were necessary for the calibration.

To 2 mL of urine, a 20 μL aliquot of IS was added. Then, 4 g of sodium chloride, 5 mL of diethyl ether and 1 mL 6 M HCL were added and samples agitated for 15 min before centrifugation at 2000 g for 5 min. The organic phase was transferred to a 15 mL glass tube. The aqueous phase was re-extracted with 5 mL ethyl acetate and agitation for 15 min before centrifugation at 2000 g for 5 min. The organic phase from this second extraction was combined with the first organic phase before evaporation to dryness under nitrogen flow. The dried sample was taken up into 1 mL of (50/50) methanol and 2 mM ammonium formiate (pH 3) and 5 μL injected into the LC-MS/MS system. The calibration standards were treated in the same way after spiking of the appropriate volume of the working solutions.

The LC-MS/MS system included a Shimadzu NEXERA X2 series and a 8060 triple quadrupole mass spectrometer. Chromatographic separations were performed on an INERTSIL ODS3 column (100 × 2.10 mm, 5 μm particles) (GL Sciences INC., JAPAN). Mobile phase A contained 2 mM ammonium formiate (pH 3) and phase B included (90/10) acetonitrile and 2 mM ammonium formiate (pH 3). Identification and quantification of DMP, DMTP, DMDTP, DEP, DETP and DEDTP were performed in negative mode using MRM of a quantifier ion (125.4/63.1, 141.3/126.1, 157.3/112.1, 153.4/79.1, 169.4/95.1 and 185.3/157.2, respectively) and an additional qualifier ion (125.4/79.1, 141.3/96.1, 157.3/142.1, 153.4/125.1, 169.4/141.1 and 185.3/111.1, respectively).

### Dithiocarbamate biomonitoring in urine samples

Dithiocarbamates are measured by carbon disulfide (CS_2_) in urine after acid hydrolysis at high temperature. The CS_2_ produced was injected into a headspace GC/MS system. For this, Benzene ^6^D (LGC, UK) used as an IS at 2 g/L was diluted in methanol to obtain a working solution of 1 mg/L. Carbon disulfide was purchased as a solution at 100 mg/L (LGC, UK), was diluted in methanol to obtain working solutions at increasing concentrations ranging from 0.2 to 10 mg/L. These standard solutions were used to spike dithiocarbamate-free urine for the preparation of the calibration curve standards. Five calibration standards between the higher LOQ and the lower LOQ (namely, between 10 to 500 μg/L) were necessary for the calibration.

SnCl_2_ purchased from PROLABO (France), was dissolved in 5 M HCl to obtain a working solution of 5 g/L with this solution being used for the acid hydrolysis. To 2 mL samples of urine, 100 μL of IS and 3 mL of the SnCl_2_ reagent were added in headspace glass tubes and were crimped. Samples were agitated for a few second and then incubated for 15 min at 100 °C. The samples were then injected into the HS-GC-MS system.

The HS-GC-MS system included a Perkin Elmer TurboMatrix HS 40 and a Shimadzu QP 2010 quadripole mass spectrometer. Chromatographic separations were performed on a RTX1 column (30 m × 0.32 mm × 4 μm) (RESTEK, France). Carrier gas was helium. For separation, temperature was increased from 50 °C to 200 °C in 9 min. Identification and quantification of carbon disulfide were performed in impact electronic ionization mode using SIM of a quantifier ion (75.9) and a additional qualifier ion (77.9).

### Faecal metabolomics

Metabolon Inc. (Durham, NC, USA) was contracted to conduct the metabolomics analysis for human faecal samples as previously described (Mesnage et al. 2021). In brief, samples were prepared using the automated MicroLab STAR® system from Hamilton Company. In order to identify metabolites with different physicochemical properties, each sample extract was analysed on four independent instrument platforms: two different separate reverse phase ultra-high performance liquid chromatography-tandem mass spectroscopy analysis (RP/UPLC-MS/MS) with positive ion mode electrospray ionisation (ESI), a RP/UPLC-MS/MS with negative ion mode ESI, as well as a by hydrophilic-interaction chromatography (HILIC)/UPLC-MS/MS with negative ion mode ESI. The details of the solvents and chromatography used are as described (Ford et al. 2020).

Raw data was extracted, peak-identified and QC processed using Metabolon’s hardware and software as previously described (DeHaven et al. 2010). Faecal metabolites were identified by comparison to libraries of authenticated standards with known retention time/index, mass to charge ratio, chromatographic and MS/MS spectral data. Peak area values allow the determination of relative quantification among samples (Evans et al. 2009).

### Faecal Shotgun metagenomics

Faecal samples were collected at home by the recruited volunteers and stored at −80[°C at King’s College London. DNA was extracted from 100 mg faecal samples using the Quick-DNA Fecal/Soil Microbe Miniprep Kit (ZymoResearch) according to the manufacturer’s instructions. Minor adaptations were made as previously described (Ducarmon et al. 2020) as follows: 1. bead beating was performed at 5.5 m/s for three times 60 seconds (Precellys 24 homogeniser, Bertin Instruments) and 25 μl elution buffer was used to elute the DNA, following which the eluate was run over the column once more to increase DNA yield. A negative control (no sample added) and a positive control (ZymoBIOMICS Microbial Community Standard, ZymoResearch) were processed for DNA extraction and subsequently sequenced. DNA was quantified using the Qubit HS dsDNA Assay kit on a Qubit 4 fluorometer (Thermo Fisher Scientific).

Shotgun metagenomics was performed under contract by GenomeScan (Leiden, The Netherlands). The NEBNext® Ultra II FS DNA module (cat# NEB #E7810S/L) and the NEBNext® Ultra II Ligation module (cat# NEB #E7595S/L) were used to process the samples. Fragmentation, A-tailing and ligation of sequencing adapters of the resulting product was performed according to the procedure described in the NEBNext Ultra II FS DNA module and NEBNext Ultra II Ligation module Instruction Manual. The quality and yield after sample preparation was measured with the Fragment Analyzer. The size of the resulting product was consistent with the expected size of approximately 500-700 bp. Clustering and DNA sequencing using the NovaSeq6000 was performed according to manufacturer’s protocols. A concentration of 1.1 nM of DNA was used. DNA sequencing data was acquired using NovaSeq control software NCS v1.6.

Shotgun metagenomics datasets were analysed with Rosalind, the BRC/King’s College London high-performance computing cluster. First, data was pre-processed using the software package pre-processing v0.2.2 (https://anaconda.org/fasnicar/preprocessing). In brief, this package concatenates all forward reads into one file and all reverse reads into another file; then uses trim_galore to remove Illumina adapters, trim low-quality positions and unknown positions (UN) and discard low-quality (quality <20 or >2 Ns) or too-short reads (< 75bp); removes contaminants (phiX and human genome sequences); and ultimately sorts and splits the reads into R1, R2, and UN sets of reads. The human faecal microbiome were analysed using MetaPhlan2 (v 2.9)(Truong et al. 2015) and Humann2 (v 0.10)(Franzosa et al. 2018) with the UniRef90 database to characterise the composition and function of the faecal microbiome samples.

### Urine metabolomics

The urine metabolomics analysis is an adaptation of a protocol originally published by Tanaka and colleagues (Tanaka et al. 1980). Briefly, a liquid-liquid extraction was first performed to extract the urine organic acids after mixing the sample with 2-ketocaproic and tropic acids as internal standards (both from Sigma Aldrich (St. Louis, MO, USA)). Hydroxylamine hydrochloride (Sigma Aldrich) was added to oxidise 2-keto acids. N,O,-bis-(trimethylsilyl) trifluoroacetamide (Supelco Bellefonte, PA, USA) containing 1% trimethylchlorosilane (Supelco Bellefonte) was then added to convert organic acids to corresponding trimethylsilyl (TMS) ethers, required to impart volatility. Volatile TMS esters were separated by gas-chromatography. Detection is performed using an electron impact mass spectrometer in scan mode with a mass range between 50 and 550 m/z. Obtained spectra are compared with published spectra for the compounds of interest to achieve identification. The absolute quantification of organic acids is performed using the calibration curves of standard compounds to internal standard ratios. Concentrations were normalized to creatinine. The quality assurance of the Organic acids’ methodology was assessed by participation in the quality control scheme of the European Research Network for Diagnosis of Inherited disorders of Metabolism (ERNDIM): Qualitative urine Organic acids and Quantitative urine Organic acids (Peters et al. 2008). Precision, linearity and recovery for this method has been published (Tsoukalas et al. 2019).

### Statistical analysis

Pesticide biomonitoring data are often left-censored because a proportion of the individual’s urinary concentrations are below the level of detection. Summary statistics for pesticide urinary concentrations were calculated using a maximum likelihood estimation with R package NADA v1.6-1 (Shoari and Dube 2018) when the number of left-censored values was below 50%. In case the number of missing values (table 1) was too high (detection frequency < 20%), only the detection frequency was reported.

Random Forest classification of the 124 urine samples in which glyphosate could be measured was performed by using faecal microbiome parameters as predictors using R package Caret (version 6.0-84)(Kuhn 2008). Since the two classes were not balanced (58 non-organic food consumers and 66 organic food consumers), down-sampling was done prior to processing with the *trainControl* function. Input variables were scaled and centred. The optimisation of the number of variables for splitting at each tree node (mtry) was done with default parameters. Accuracy was estimated using repeated cross-validation (5-fold, repeated 10 times). The model was trained using 66% of the dataset while the quality of this model was evaluated using predicted sample classification of the remaining 34% of the dataset. The quality control metrics were calculated using the *confusionMatrix* function from Caret. This function calculates the overall accuracy along a 95% confidence interval, with statistical significance of this accuracy evaluated with a one-side test comparing the experimental accuracy to the ‘no information rate’.

While pesticides with a detection frequency over 80% were considered as continuous variables, those detected in 50-80% of the samples were dichotomized as detected/undetected as recommended by the European Human Biomonitoring Initiative (HMB4EU) and previously described (Harel et al. 2014). Pesticides with detection frequencies below 20% were not carried forward in the association study (table 1). The ExpWAS was conducted with a linear-mixed model considering age as a covariate and family relationship as a random effect with MaAsLin (Microbiome Multivariable Association with Linear Models) 2.0 (package version 0.99.12) (Mallick et al. 2021). Metagenome taxa detected in less than 20% of the individuals were removed and 211 species were carried forward for the association analysis. The metabolome data was log-transformed, while the metagenome taxonomic composition was transformed using an arcsine square root transformation. The Benjamini–Hochberg method was used to control the False Discovery Rate (FDR) of the MaAsLin analysis. Shannon and Simpson diversity indices, and species richness, were calculated with the vegan R package version 2.5-6. (Oksanen et al. 2019). Nonmetric multidimensional scaling of Bray-Curtis dissimilarity with stable solution from random starts, with axis scaling, was performed with vegan R package (Oksanen et al. 2019). Statistical significance of Bray-Curtis dissimilarity differences according to pesticide residue levels was calculated by Permutational Multivariate Analysis Of Variance (PERMANOVA) with 1,000 permutations (Anderson 2001).

## Acknowledgements

This work was funded by the Sustainable Food Alliance (USA) whose support is gratefully acknowledged.

## SUPPLEMENTARY MATERIAL

**Table S1.** Responses to and corresponding weights applied to question “Please indicate to what extent you consume, when available, organic fruits and vegetables?” to estimate a proxy for potential for pesticide exposure

| Response | Weight applied to fruit and vegetables (g/week) |
| --- | --- |
| 0 - I don't eat fruits and vegerables | Removed from analysis |
| 1 Almost all non-organic fruits and vegetables | 2 |
| 2 A mixture of organic and non-organic fruits and vegetables | 1 |
| 3 Almost all organic fruits and vegetables | 0.2 |

**Table S2.**
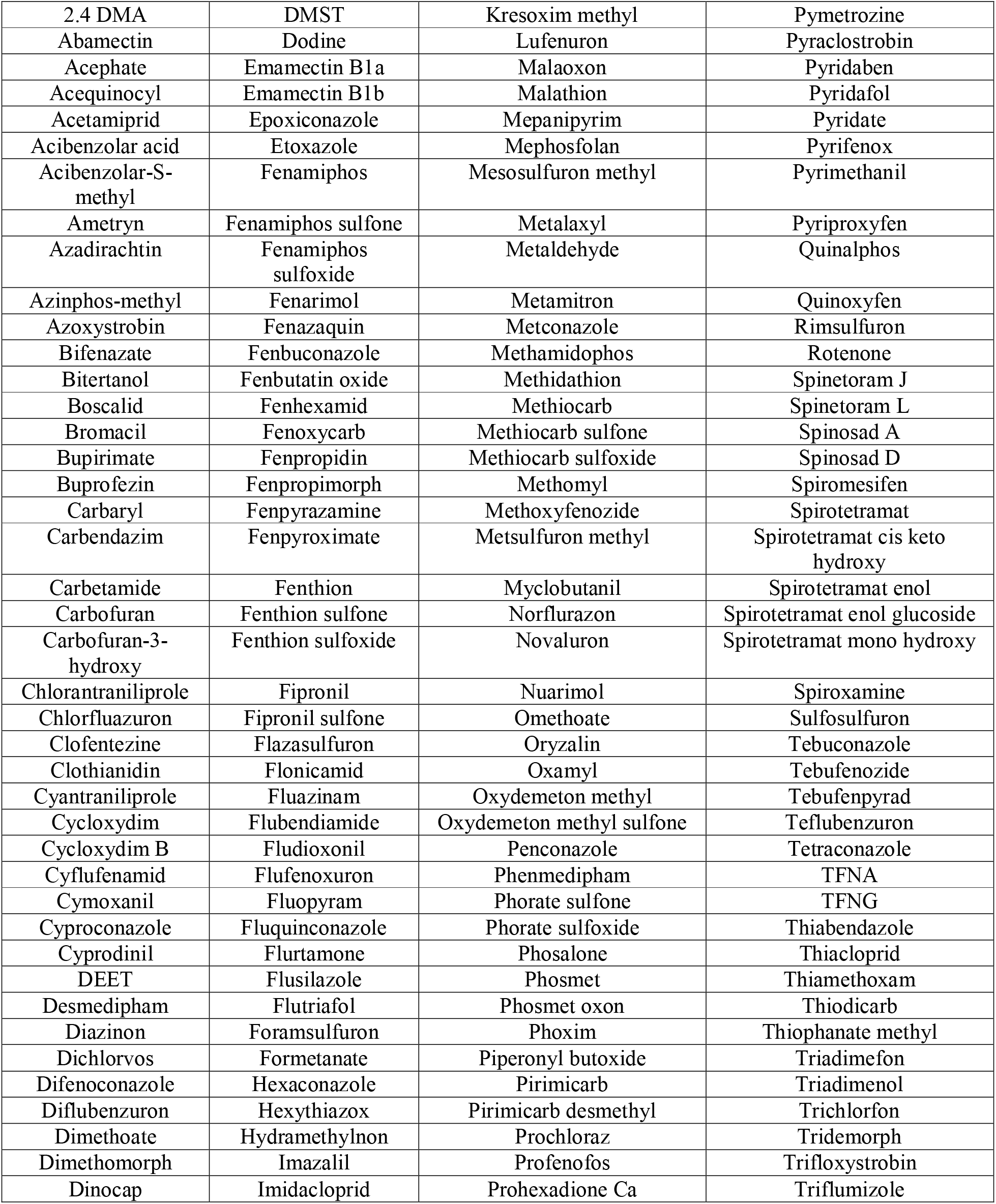
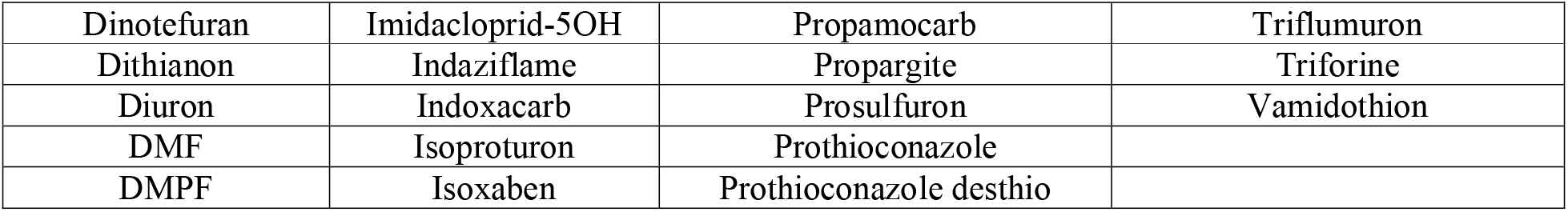
Pesticide residues (parent compounds or metabolites) which were included in the screening assay (limit of detection < 0.1 μg/L). A total of 186 compounds were measured in the urine samples from 130 twins from the TwinsUK cohort. Organophosphates, pyrethroids, and glyphosate do not appear in this table because they were measured using targeted quantitative assays.

**Table S3.** We also calculated the total exposure to organophosphates for DEs (molar sum of DEP + DETP + DEDTP), DMs (molar sum of DMP + DMTP + DMDTP), and the total DAPs (molar sum of all OP metabolites). The summary statistics values (nM) were estimated using the maximum likelihood inference for left-censored values. DF, detection frequency; IQR, 5 percent and 95 percent quantiles.

| Metabolites | LOD | DF | median | max | IQR |
| --- | --- | --- | --- | --- | --- |
| sum DMP | - | 60 | 42.7 | 762 | 5.9 – 311.1 |
| sum DEP | - | 74.6 | 16.4 | 1202 | 2.4 – 113.4 |
| sum DAP | - | 91.5 | 34.3 | 1202 | 3.0 – 381.4 |

**Table S4.** Summary of the most significant associations between urinary excretion of insecticide residues and the composition of the faecal microbiota. Associations with FDR < 0.2 for both creatinine adjusted and unadjusted models are reported.

| response | predictor | creatinine adjusted |  |  |  | unadjusted |  |  |  |
| --- | --- | --- | --- | --- | --- | --- | --- | --- | --- |
|  |  | coef | sd | pval | qval | coef | sd | pval | qval |
| metagenomics |  |  |  |  |  |  |  |  |  |
| *Olsenella scatoligenes* | Cis.Cl2CA | 0.083 | 0.011 | 1.1E-09 | 2.2E-07 | 0.071 | 0.011 | 4.7E-08 | 2.0E-05 |
| *Olsenella scatoligenes* | Trans.Cl2CA | 0.077 | 0.014 | 1.2E-06 | 7.8E-04 | 0.064 | 0.013 | 1.7E-05 | 7.1E-03 |
| *Collinsella stercoris* | sumDAP | 0.028 | 0.008 | 3.6E-04 | 1.7E-01 | 0.026 | 0.007 | 2.3E-04 | 4.9E-02 |
| *Clostridium symbiosum* | X3.PBA | 0.090 | 0.018 | 7.3E-05 | 3.3E-02 | 0.065 | 0.015 | 2.4E-04 | 5.0E-02 |
| *Streptococcus cristatus* | X3.PBA | -0.012 | 0.003 | 1.6E-04 | 3.3E-02 | -0.015 | 0.000 | 5.3E-39 | 2.2E-36 |
| metabolomics |  |  |  |  |  |  |  |  |  |
| N-succinyl-isoleucine | Br2CA | -0.116 | 0.035 | 1.2E-03 | 1.3E-01 | -0.089 | 0.031 | 4.5E-03 | 1.9E-01 |
| oleoyl-linoleoyl-glycerol | Trans.Cl2CA | 0.234 | 0.071 | 1.3E-03 | 1.5E-01 | 0.193 | 0.064 | 3.3E-03 | 1.4E-01 |
| oleoyl-GPC | Trans.Cl2CA | 0.248 | 0.082 | 3.1E-03 | 1.9E-01 | 0.258 | 0.073 | 6.2E-04 | 6.9E-02 |

**Table S5.** We calculated if the alpha and beta diversity could be influenced by pesticide levels. The p-values for differences in beta diversity were estimated with a PERMANOVA test conducted with R-vegan function adonis. P-values for different alpha diversity indices (p-shannon, Shannon index; p-simpson, Simpson index, p-observed; observed species) were calculated with linear-mixed models considering considering age as a covariate and the family relationship as a random effect.

| **Metabolites** | **p_beta** | **p_shannon** | **p_simpson** | **p_observed** |
| --- | --- | --- | --- | --- |
| Glyphosate | 0.17 | 0.13 | 0.45 | 0.02 |
| Trans.Cl2CA | 0.35 | 0.73 | 0.29 | 0.95 |
| Cis.Cl2CA | 0.2 | 0.70 | 0.78 | 0.81 |
| 3-PBA | 0.05 | 0.43 | 0.90 | 0.26 |
| DEP | 0.13 | 0.43 | 0.66 | 0.04 |
| Br2CA | 0.09 | 0.75 | 0.52 | 0.29 |
| sumDMP | 0.05 | 0.59 | 0.85 | 0.27 |
| sumDEP | 0.11 | 0.44 | 0.67 | 0.04 |
| sumDAP | 0.05 | 0.96 | 0.93 | 0.31 |

**Table S6.** Comparative organic acids analysis in urine between a group of 28 individuals who did not present detectable glyphosate levels in their urine compared to a group of 33 individuals with detectable glyphosate levels. Concentrations of organic acids are expressed as mmol/mol Creatinine. Mean ± are provided along p-values of a t-test adjusted using the FDR method.

|  | **Undetected** | **Detected** | **Adjusted p-value** |
| --- | --- | --- | --- |
| **citric acid** | 150 ± 110 | 170 ± 120 | 0.92 |
| **aconitic acid** | 32 ± 22 | 44 ± 31 | 0.92 |
| **isocitric acid** | 6.8 ± 4.6 | 8.0 ± 5.2 | 0.92 |
| **2-ketoglutaric acid** | 8.2 ± 10 | 8.1 ± 6.4 | 0.98 |
| **succinic acid** | 12 ± 23 | 13 ± 19 | 0.98 |
| **fumaric acid** | 0.046 ± 0.25 | 0.15 ± 0.48 | 0.92 |
| **malic acid** | 1.1 ± 0.51 | 1.2 ± 1.4 | 0.92 |
| **3-hydroxy-3-methylglutaric acid** | 0.082 ± 0.30 | 0.21 ± 0.45 | 0.92 |
| **lactic acid** | 71 ± 170 | 42 ± 53 | 0.92 |
| **pyruvate acid** | 6.2 ± 4.1 | 6.6 ± 6.7 | 0.98 |
| **3-hydroxybutyric acid** | 1.2 ± 0.78 | 1.4 ± 1.4 | 0.92 |
| **pyroglutamic acid** | 16 ± 9.1 | 18 ± 11 | 0.92 |
| **3-hydroxyisovaleric acid** | 7.4 ± 5.0 | 10 ± 7.2 | 0.92 |
| **methylmalonic acid** | 1.1 ± 0.48 | 1.1 ± 0.54 | 0.98 |
| **homovannilic acid** | 1.9 ± 1.1 | 2.2 ± 1.2 | 0.92 |
| **5-hydroxyindoloacetic acid** | 2.2 ± 1.3 | 3.5 ± 4.5 | 0.92 |
| **vanillilmandelic acid** | 1.5 ± 0.66 | 1.8 ± 0.96 | 0.92 |
| **4-hydroxyphenylacetic acid** | 10 ± 11 | 12 ± 5.5 | 0.92 |
| **orotic acid** | 0.050 ± 0.26 | 0.030 ± 0.17 | 0.97 |
| **glutaric acid** | 0.97 ± 0.40 | 0.92 ± 0.30 | 0.92 |
| **2-hydroxyglutaric acid** | 4.2 ± 2.5 | 3.7 ± 2.2 | 0.92 |
| **glycolic acid** | 44 ± 28 | 42 ± 26 | 0.97 |
| **oxalic acid** | 7.7 ± 5.4 | 9.2 ± 10 | 0.92 |
| **glyceric acid** | 1.9 ± 1.1 | 2.1 ± 1.7 | 0.92 |
| **2-hydroxyisobutyric acid** | 13 ± 6.7 | 13 ± 7.3 | 0.98 |
| **X2.hydroxybutyric acid** | 0.36 ± 0.61 | 0.54 ± 0.80 | 0.92 |
| **ethylmalonic acid** | 1.5 ± 0.72 | 1.5 ± 0.71 | 0.98 |
| **methylsuccinic acid** | 0.046 ± 0.25 | 0.15 ± 0.84 | 0.92 |
| **adipic acid** | 1.5 ± 0.89 | 1.6 ± 0.91 | 0.92 |
| **suberic acid** | 0.94 ± 0.64 | 0.94 ± 0.68 | 0.98 |
| **methylcitric acid** | 0.71 ± 0.53 | 0.71 ± 0.48 | 0.98 |
| **4-hydroxyphenypyruvic acid** | 1.1 ± 0.41 | 1.0 ± 0.41 | 0.92 |

**Figure S1.**
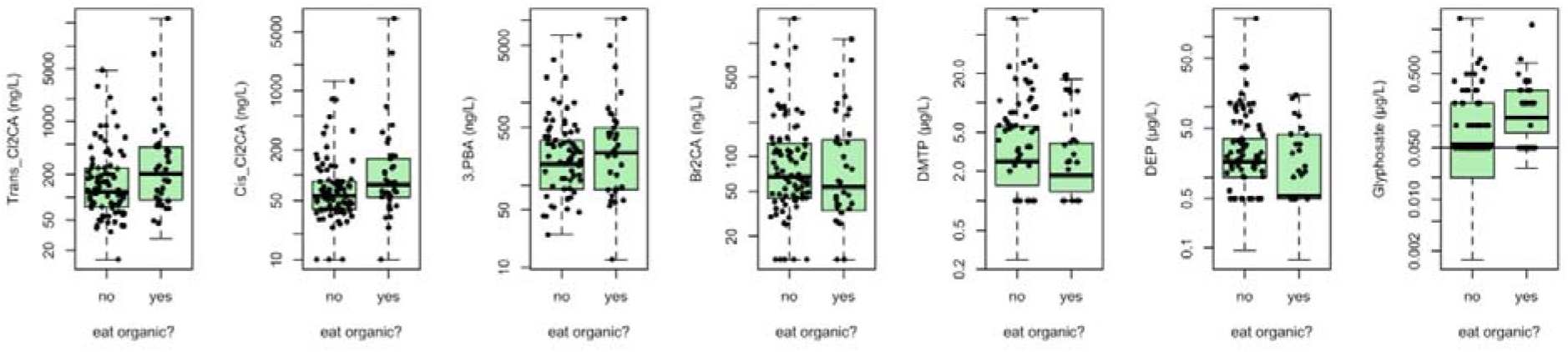
Urinary pesticide excretion between individuals eating almost all organic, or a mixture of organic and non-organic fruits and vegetables.

**Figure S2.**
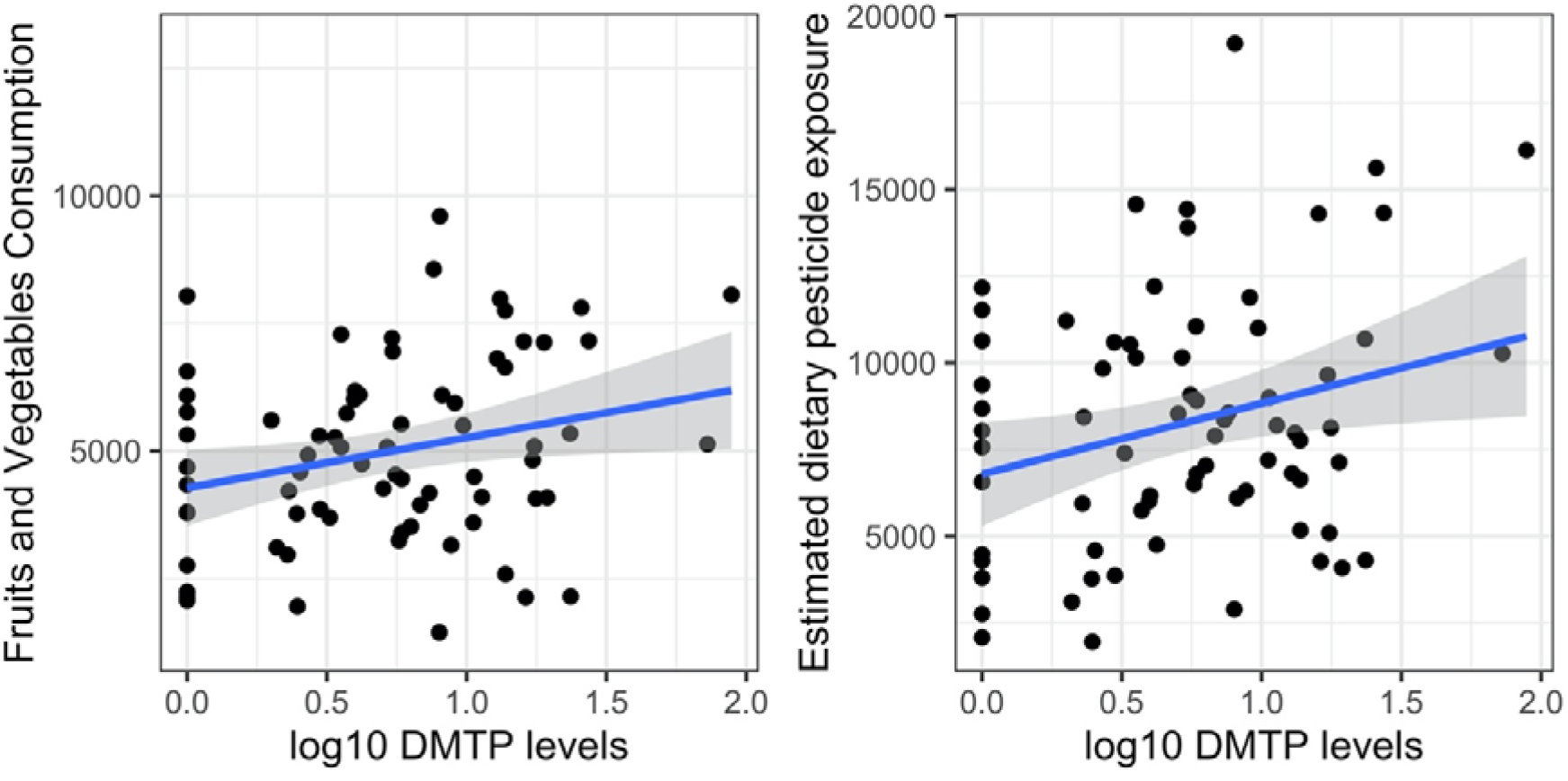
Urinary concentrations in DMTP levels correlate with the estimate dietary pesticide exposure.

**Figure S3.**
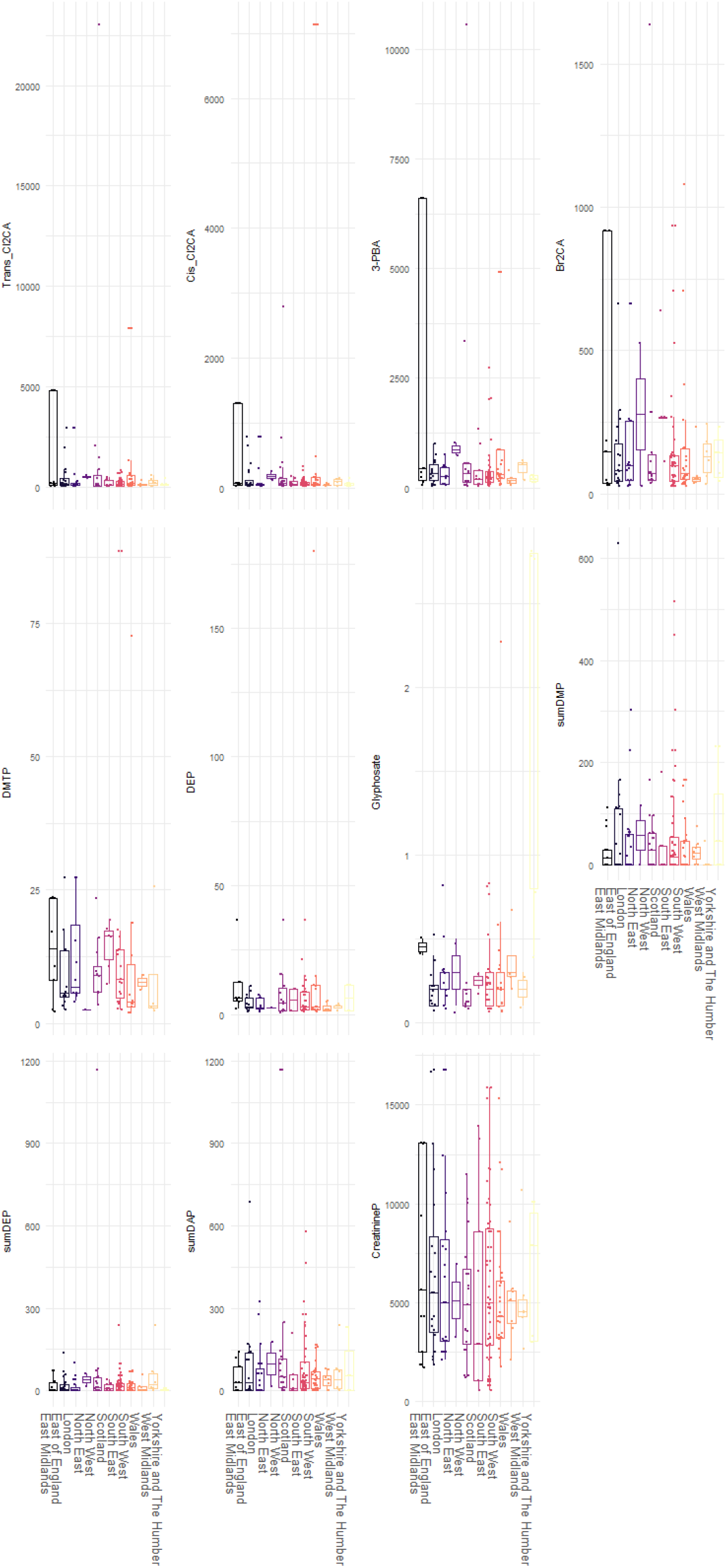
Pesticide urinary excretion across 9 UK regions.

**Figure S4.**
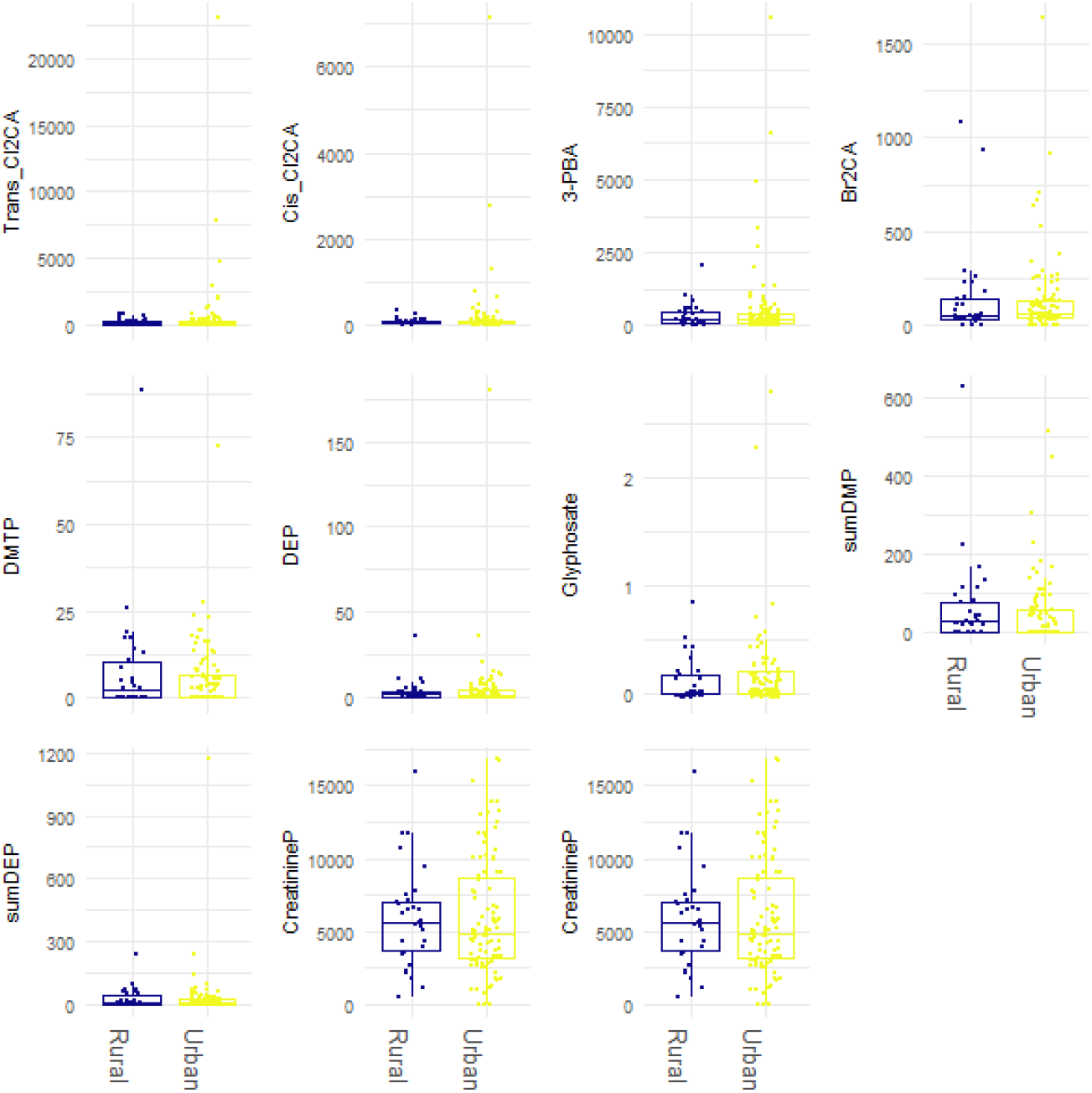
Rural/urban classification had no influence on pesticide urinary excretion.

**Figure S5.**
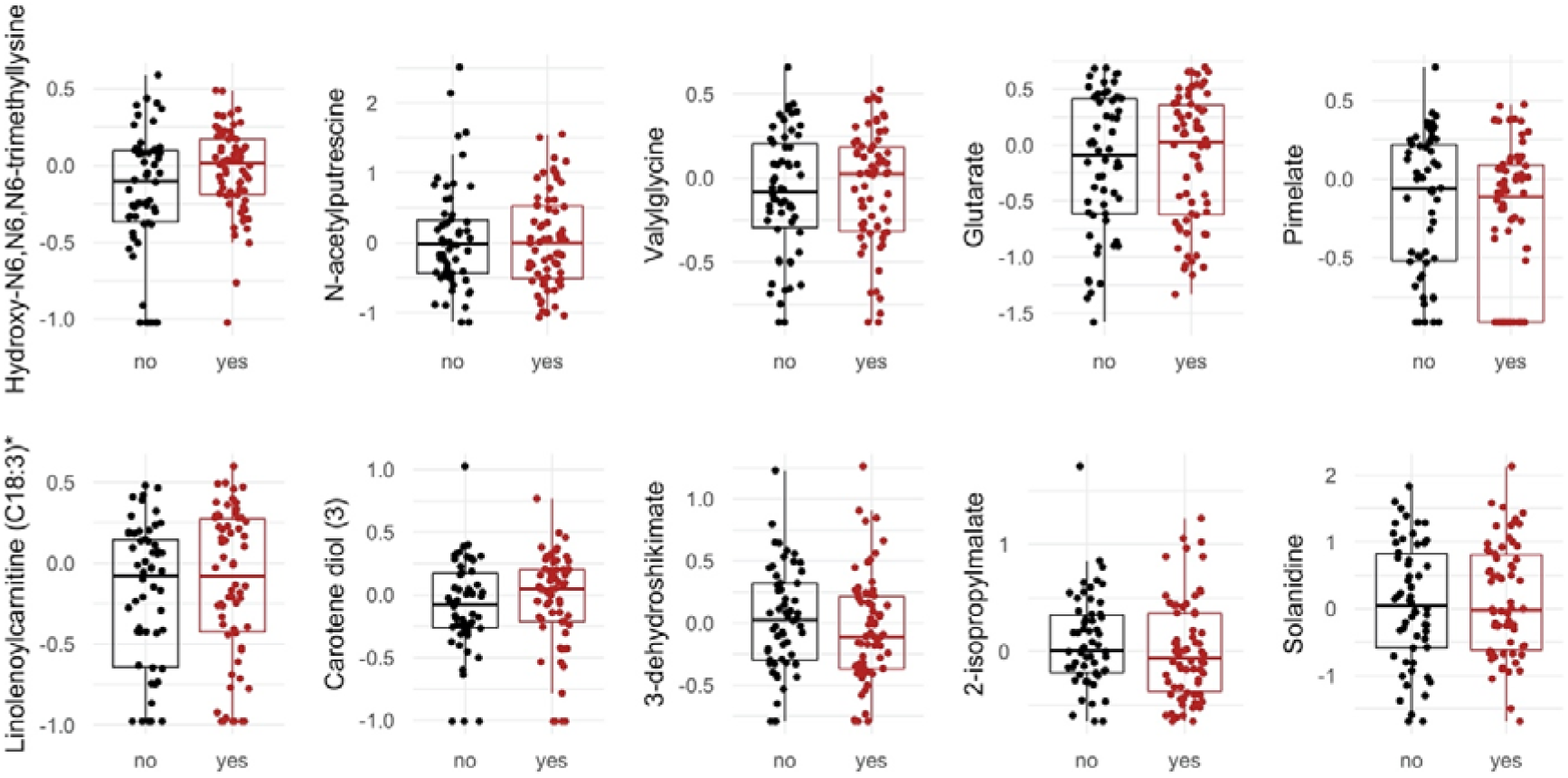
The 10 metabolites which are known to characterise the effects of glyphosate on the faecal metabolome in rats. Log-transformed abundance values are shown as box plots.

**Figure S6.**
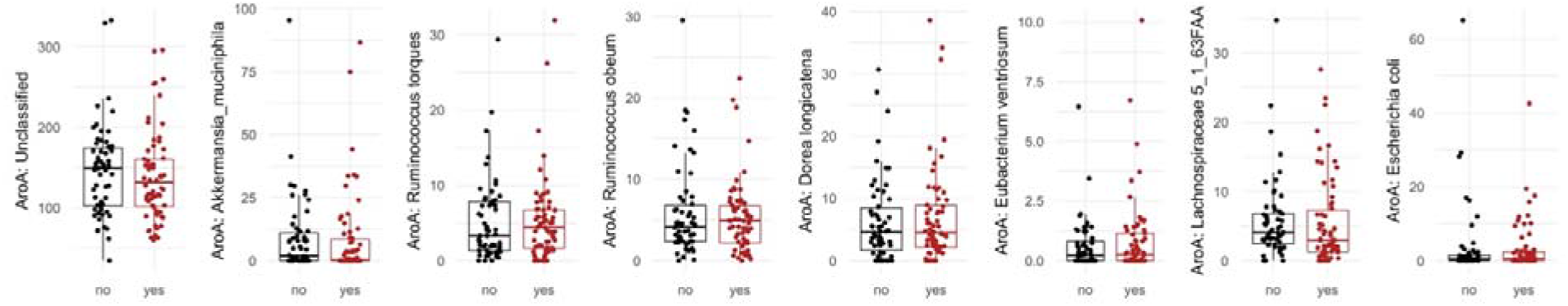
Abundance of genes from the shikimate pathway among bacteria which are identified as major contributors to the total abundance of this pathway. Log-transformed abundance values are sho

